# Wt1 positive neurons in the hindbrain are essential for respiration

**DOI:** 10.1101/2019.12.20.884361

**Authors:** Danny Schnerwitzki, Christian Hayn, Birgit Perner, Christoph Englert

## Abstract

Neuronal networks commonly referred to as central pattern generator (CPG) networks coordinate the generation of rhythmic activity like locomotion and respiration. These networks are proposed to exhibit a high degree of homology in their development. Their establishment is influenced by a variety of transcription factors. One of them is the Wilms tumor protein Wt1 that is present in dI6 neurons of the ventral spinal cord, which are involved in the coordination of locomotion. Here we report about the so far undescribed presence of Wt1 in neurons of the caudoventral medulla oblongata and their impact on respiration. By performing marker analyses, we were able to characterize these Wt1 positive (+) cells as dB4 neurons. The temporal pattern of Wt1 occurrence suggests a role for Wt1 in the differentiation of dB4 neurons during embryonic and postnatal development. Conditional knockout of *Wt1* in these cells caused an altered population size of V0 neurons already in the developing hindbrain leading to a decline in the respiration rate in the adults. Thereby, we confirmed and extended the so far proposed homology between neurons of the dB4 domain in the hindbrain and dI6 neurons of the spinal cord in terms of development and function. Ablation of Wt1+ dB4 neurons resulted in the death of neonates due to the inability to initiate respiration suggesting a vital role for Wt1+ dB4 neurons in breathing. These results extend the role of Wt1 in the CNS and show that in addition to its function in differentiation of dI6 neurons it also contributes to the development of dB4 neurons in the hindbrain that are critically involved in the regulation of respiration.

## Introduction

In vertebrates, the generation of rhythmic activity (*e.g.* breathing, chewing and walking) is mediated by a network of neurons, commonly referred to as central pattern generator (CPG) networks (Hooper, 2000). These networks are responsible to generate repetitive patterns of motor activity that do not require sensory input. However, sensory input is crucial for the refinement of the CPG activity in response to external events.

In 1914, T. G. Brown reported the first observation of intrinsic networks in the spinal cord being capable of generating motor activity in a rhythmic fashion (Brown, 1914). Further investigations revealed different types of CPGs that also vary in their complexity. CPGs responsible for protective reflexes such as coughing and swallowing are almost completely autonomous (Ota et al., 2013). Other CPGs such as the ones that coordinate breathing are continuously active and are modulated by changes in the metabolism, *e.g.* CO_2_ concentration in the blood (Hernandez-Miranda and Birchmeier, 2015). Finally, there are more complex circuits like the locomotor CPGs in the spinal cord. These CPGs are activated and controlled by command centers that are located in the brain and are refined by sensory feedback from the muscles that they activate (Grillner and Jessell, 2009).

Respiration is the inhalation of air from the outside into the lungs (inspiration) and exhalation of air in the opposite direction (expiration), which is achieved by the rhythmical movement of intercostal muscles and diaphragm. Under resting conditions, inspiration is driven actively by neurons and muscles, whereas expiration is based on passive relaxation. During increased demand of oxygen, e.g. due to physical activity or excitement, expiration becomes actively driven as well. The breathing movements are coordinated by the respiratory CPG in the brainstem (Fig. 1A). The generation and regulation of the breathing rhythm is achieved by neurons concentrated in three main brainstem areas (Alheid GF, 2008): the pontine respiratory group (PRG) within the dorsolateral pons, the dorsal respiratory group within the nucleus of the solitary tract (NTS) and the ventrolateral medulla from the level of the spinal-medullary junction through the level of the facial nucleus, which is named ventral respiratory column (VRC). The VRC contains the major neuron populations that build up the respiratory CPG. These neurons are clustered in different compartments. The excitatory pacemaker neurons, which are primarily responsible for rhythmic inspiration, are localized in the pre-Bötzinger complex (McKay et al., 2005). The pre-Bötzinger complexes can be found in each hemisphere of the hindbrain and are bilaterally connected. This allows the generation of a synchronous respiratory rhythm that is needed to innervate the muscles of the thorax simultaneously in each half of the body (Bouvier et al., 2010). Another compartment involved in the regulation of respiration is the Bötzinger complex. It mainly contains neurons that inhibit inspiration and therefore allow passive expiration during normal breathing (Ezure et al., 2003). The rostral and caudal ventral respiratory group (rVRG and cVRG) encompass premotor neurons that are activated by neurons from the pre-Bötzinger complex and inhibited by neurons from the Bötzinger complex (Alheid and McCrimmon, 2008). These premotor neurons of the rVRG and cVRG transmit information directly to phrenic motor neurons in the cervical spinal cord, which ultimately innervate the muscles of the diaphragm and the intercostal muscles and lead to inspiration.

**Fig. 1.**
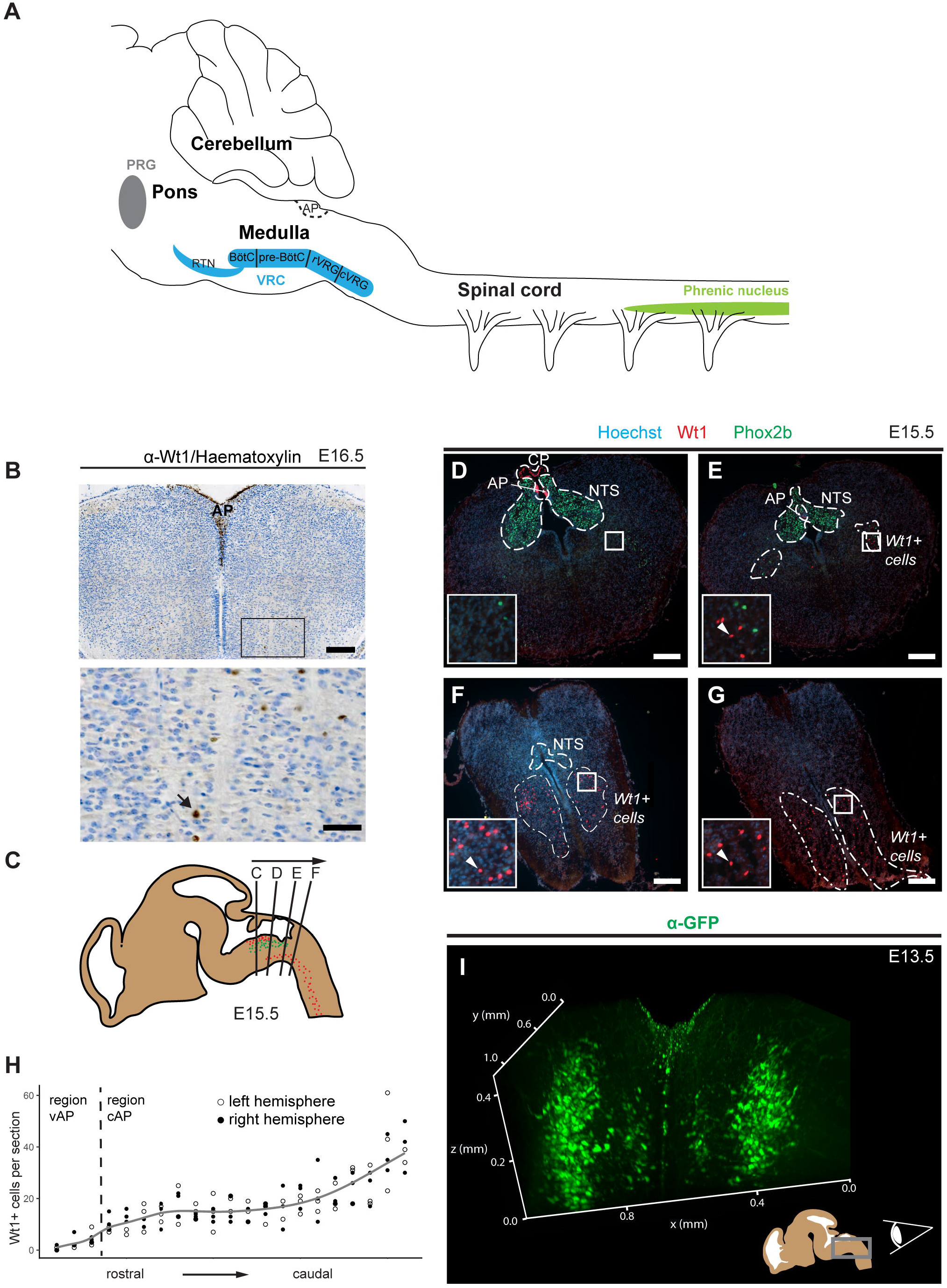
Novel domain of Wt1+ cells in the caudoventral medulla. **(A)** The respiratory center is situated in the pons and the medulla of the mouse brain. It consists of the pontine respiratory group (PRG) and the ventral respiratory column (VRC). The VRC contains the neurons that comprise the respiratory CPG. These neurons are clustered in different compartments such as the retrotrapezoid nucleus (RTN), the Bötzinger-(BötC) and the pre-Bötzinger complex (pre-BötC) as well as the rostral and caudal ventral respiratory group (rVRG and cVRG). BötC and pre-BötC contain neurons that are responsible for expiration and inspiration, respectively. These neurons are modulated by the RTN that possesses sensory function for CO_2_. For inspiration, neurons in the pre-BötC innervate neurons in the rVRG and the cVRG that activate motor neurons in the phrenic nucleus of the cervical spinal cord. These motor neurons finally innervate the muscles of the diaphragm leading to their contraction and thereby inspiration. AP: area postrema. **(B)** Immunohistochemical analysis of Wt1 on sections of the hindbrain from E16.5 embryos reveals a region of the caudoventral medulla where *Wt1* expressing cells occur. This region is ventral to the V-shaped area postrema (AP) where non-neuronal Wt1+ cells can be found. Higher magnification image shows area in the ventral medulla where Wt1+ cells can be found. Nuclei where counterstained with haematoxylin, which show that Wt1 localizes in the nucleus (black arrow). Orientation of the images: dorsal-up and ventral-down. Scale bar top row: 200 μm; bottom row: 50 μm. **(C)** The scheme represents a sagittal brain view demonstrating the levels of the coronal sections B, C, D and E. Cells harboring Wt1 are depicted as red dots, those harboring Phox2b are green. **(D)** In coronal section from rostral levels of the medulla oblongata, the localization of Wt1 is restricted to the choroid plexus (CP) and the area postrema (AP). **(E)** At a particular position ventral of the AP where the CP is no longer observable, another population of Wt1+ cells occur in the ventral proportion of both hemispheres. **(F, G)** The area that the Wt1 populations in both hemispheres extends and enlarges in caudal direction. Sections were obtained from embryos staged E15.5. Scale bars: 200 µm. **(H)** The distribution of Wt1+ cells in coronal sections was determined along the rostrocaudal axis. The number of Wt1+ cells increases in caudal direction within the region ventral to the AP (vAP). In the region caudal to the AP (cAP), the Wt1 cell number stabilizes until it increases further caudally towards the spinal cord. Linear regression between cell number and position along the rostrocaudal axis is shown in grey. The cell number of Wt1+ cell in every 5th coronal section (12 µm) was determined. n = 3 individuals (E15.5). **(I)** 3D reconstruction of the caudal hindbrain shows the distribution of Wt1+ cells (green). To label the cells, hindbrains from E13.5 *Wt1^GFP^* embryos were used. GFP was detected using whole mount immunofluorescence staining followed by tissue clearing and light sheet microscopy. Wt1+ neurons are distributed in two lateral columns and two smaller and more medial columns. View: from caudal to rostral (as shown in the scheme).

This high grade of compartmentalization is due to a distinct spatial patterning during embryonic development. The anterior-posterior patterning is mainly driven by members of the *Hox* gene family (Philippidou P, 2013). Their gene products instruct the posterior identity of hindbrain but also of the spinal cord. Floor and roof plate act as organizers to establish dorsoventral patterning of spinal cord and hindbrain via the secretion of morphogenic signals (Jessell, 2000). A concentration gradient of these morphogenic proteins determine distinct progenitor cell domains in the ventricular zone along the dorsoventral axis. These domains give rise to particular cell populations, which differentiate and migrate to the mantle zone. The individual cell populations build up the neuronal circuits of the spinal cord and hindbrain.

The spatial patterning and the neuronal cell fate is driven by a specific set of different transcription factors that are similar between the spinal cord and the hindbrain (Hernandez-Miranda LR, 2016). The transcription factor Lbx1, for instance, is crucial for the development of dorsal class B neurons: dB1 - dB4 in the hindbrain (Sieber et al., 2007) and dI4 – dI6 neurons in the spinal cord (Gross et al., 2002; Müller et al., 2002). Thus, the inactivation of *Lbx1* alters the developmental program of somatosensory neurons to a more dorsal neuron phenotype associated with impairments in locomotion and respiration, respectively (Müller et al., 2002; Pagliardini S, 2008). Similarly, we and others have recently shown that inactivation of the Wilms tumor suppressor gene *Wt1* alters the composition of ventral neurons in the spinal cord as well as the locomotion behavior (Haque et al., 2018; Schnerwitzki et al., 2018). The Wilms tumor protein Wt1 plays an important role in the development and homeostasis of mostly mesoderm-derived tissues like gonads, kidneys, spleen and heart (Chau et al., 2011; Dong et al., 2015; Herzer et al., 1999; Kreidberg et al., 1993; Moore et al., 1999) and has only recently been shown to also have a function in the CNS. Here, it acts as a zinc finger transcription factor on the development of dI6 neurons in the spinal cord, which participate in the control of locomotion. Since a homology between dI6 neurons of the spinal cord and class B neurons of the hindbrain have been proposed in terms of development (Hernandez-Miranda LR, 2016), we aimed to elucidate whether Wt1 also occurs in class B neurons of the hindbrain and whether this cells share functional similarities with dI6 neurons of the spinal cord.

## Results

### Wt1 is expressed in cells of the caudoventral medulla

When investigating Wt1’s role during the development of dI6 neurons in the spinal cord (Schnerwitzki et al., 2018), we incidentally discovered a so far unreported expression domain of *Wt1* in the hindbrain (Fig. 1B). Immunohistochemical staining of coronal sections through the embryonic medulla confirmed the already reported occurrence of Wt1+ cells in a region below the fourth ventricle called area postrema (AP). In addition, we observed further Wt1+ cells in a more ventral region of the medulla.

To examine the distribution of the Wt1+ cells in the ventral parts of the developing medulla, consecutive coronal sections from embryonic mice at stage E15.5 were analyzed (Fig. 1C). The nucleus of the solitary tract (NTS) was labelled by Phox2b staining for orientation within the hindbrain. At the very rostral sections (Fig. 1D), Wt1+ cells were solely detected in the AP and caudal parts of the choroid plexus of the fourth ventricle but not in the ventral region of the medulla. Further caudal the dorsal occurrence of Wt1+ cells is restricted to the AP (Fig. 1E), while additional Wt1+ cells appear in two ventral areas; one per hemisphere. The extent of these areas increases in caudal direction (Fig. 1F), whereas the extent of the NTS decreases until no Phox2b+ cells remain (Fig. 1G). This is where the dimension of the Wt1+ cell population is the largest and merges with the population in the spinal cord. No discrete border was observed between both Wt1 population, the one in the hindbrain and the one in the spinal cord.

The distribution of the ventral Wt1+ cells in both hemispheres was determined along the rostrocaudal axis in more detail (Fig. 1H). To this end, Wt1+ cells were counted on consecutive coronal sections. Wt1+ cells were clustered in two regions: the rostral part of the Wt1 population is located ventral to both the AP and the NTS. The second region includes the caudal fraction of the Wt1 cell population, still ventral of the NTS but caudal of the AP. This region extends to the last section where Phox2b+ cells, as a marker for the NTS, were still detected. This restriction ensures that only Wt1+ cells of the hindbrain are included in the analysis but not those of the spinal cord. The quantification revealed that the number of Wt1+ cells in the region ventral to the AP increases in caudal direction. In the region caudal to the AP, the number of Wt1+ cells is constant over a certain area before it further increases at the transition from the hindbrain to the spinal cord. The separate quantification of Wt1+ cells between the left and right hemisphere of the medulla showed that the Wt1+ cells are evenly distributed on both sides.

Bearing in mind the limitations of extrapolating a spatial distribution on the basis of serial sections, we carried out a 3D reconstruction of the special distribution of Wt1+ cells in the ventral medulla using a *Wt1^GFP^* reporter mouse line. After whole mount immunofluorescence staining, subsequent tissue clearing and applying light sheet microscopy, the 3D reconstruction revealed two parallel columns of Wt1+ cells in the ventral area of the medulla, one in each hemisphere (Fig. 1I and Suppl. Vid. 1). Moreover, two minor columns were detected, which are situated more medial. Thus, Wt1+ cells in the ventral hindbrain appear as four domains, namely two major and more laterally located ones and two minor, more medially located ones.

### Temporal distribution of Wt1+ cells in the caudoventral hindbrain

To determine when the Wt1+ cells occur in the hindbrain for the first time, brains of developmental stages E11.5, E12.5 and E13.5 were analyzed (Fig. 2A). While until E12.5 no Wt1+ cells were detected, a first occurrence of Wt1+ cells in the ventral medulla was observed at E13.5. Wt1+ cells of the AP appear at this stage for the first time as well.

**Fig. 2.**
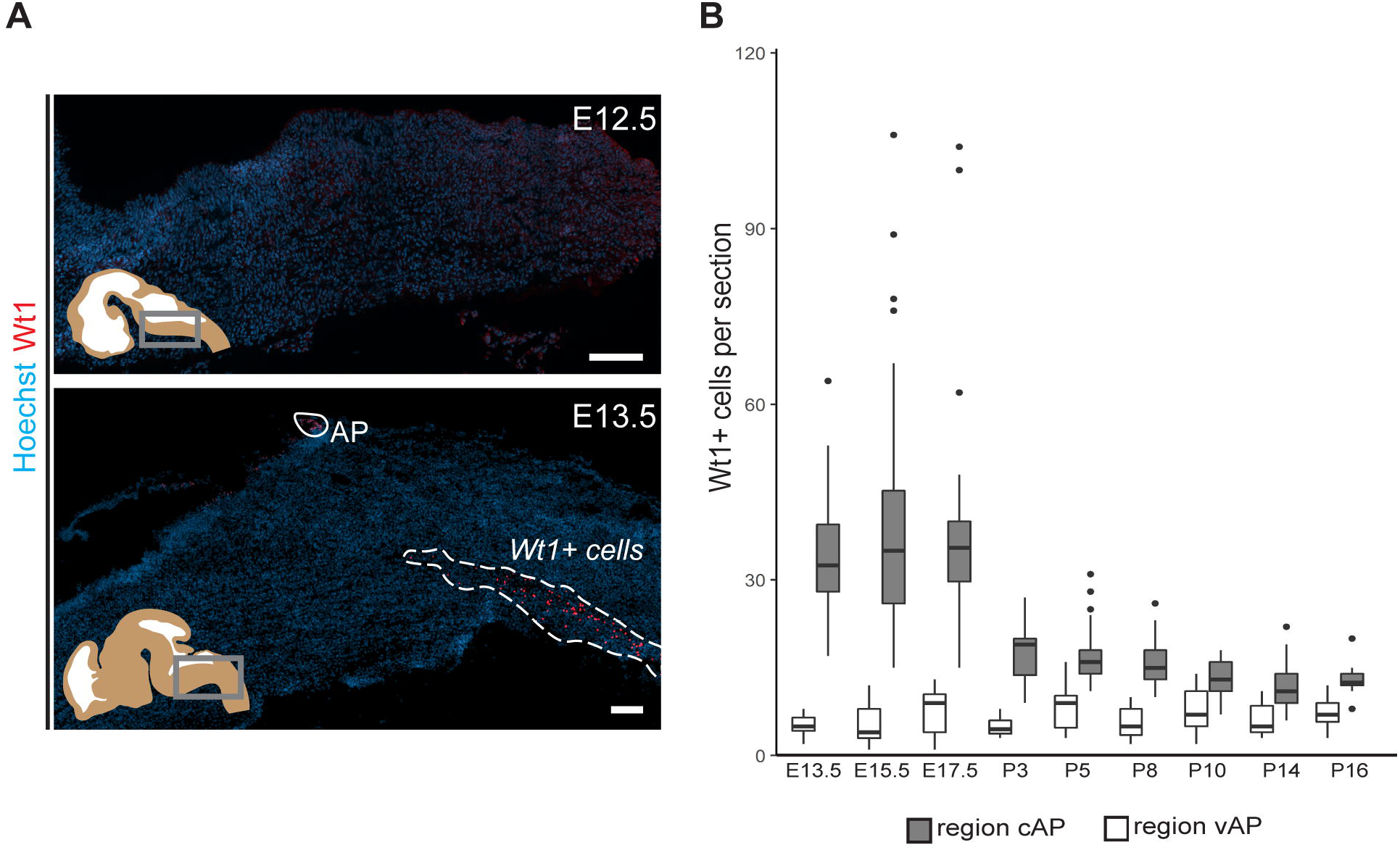
Occurrence of Wt1 in cells of the caudoventral hindbrain during development. **(A)** Immunofluorescence analyses of sagittal sections from embryonic hindbrains reveal presence of Wt1+ cells in the AP and in the caudoventral hindbrain at developmental stage E13.5. At earlier stages, no Wt1+ cells are detectable in the hindbrain. Scale bars: 100 µm. **(B)** The average number of Wt1+ cells was determined for every 5th coronal hindbrain section from different embryonal and postnatal stages. The cells were categorized by their occurrence relative to the AP in a region caudal of the AP (cAP) and ventral of the AP (vAP). The number of Wt1+ cells in the region cAP is higher than in the region vAP. Postnatally, the number of Wt1+ cells decreases in the region vAP to cell numbers comparable to the region cAP. n per stage = 2-3 idividuals

Next we investigated how the temporal distribution of Wt1+ cells in the ventral hindbrain changes during development (Fig. 2B). The average number of Wt1+ cells in coronal section in both the regions ventral and caudal to the AP was determined. The amount of cells ventral to the AP remains constant in all investigated developmental stages. In contrast, more cells occur in the region caudal to the AP at embryonal stages and the cell number also varies significantly. At the postnatal stages, however, the number of Wt1+ cells decreases significantly in this region. This points to a dynamic regulation of Wt1 in those cells during development depending on their position in the hindbrain.

### Wt1 expressing cells in the caudoventral medulla are dB4 neurons

Due to the homology between spinal cord and hindbrain in terms of development, it has been proposed that Wt1 would be a specific marker for dB4 neurons in the hindbrain as it is for dI6 neurons in the spinal cord (Hernandez-Miranda LR, 2016) (Fig. 3A). However, no primary data have been made available so far. In order to verify this hypothesis, immunofluorescence analysis was performed using hindbrain section from E16.5 embryos (Fig. 3B). Wt1+ cells in the ventral medulla were found to express the neuronal marker *NeuN* showing that these cells are neurons. Moreover, the Wt1+ neurons can also be labelled with antibodies against the transcription factors Lb×1 and Bhlhb5 that commonly occur in the most ventral Lbx1 domain that gives rise to dB4 neurons (Fig. 3A). Thus, these suggest Wt1 to be a marker for at least a subpopulation of dB4 neurons.

**Fig. 3.**
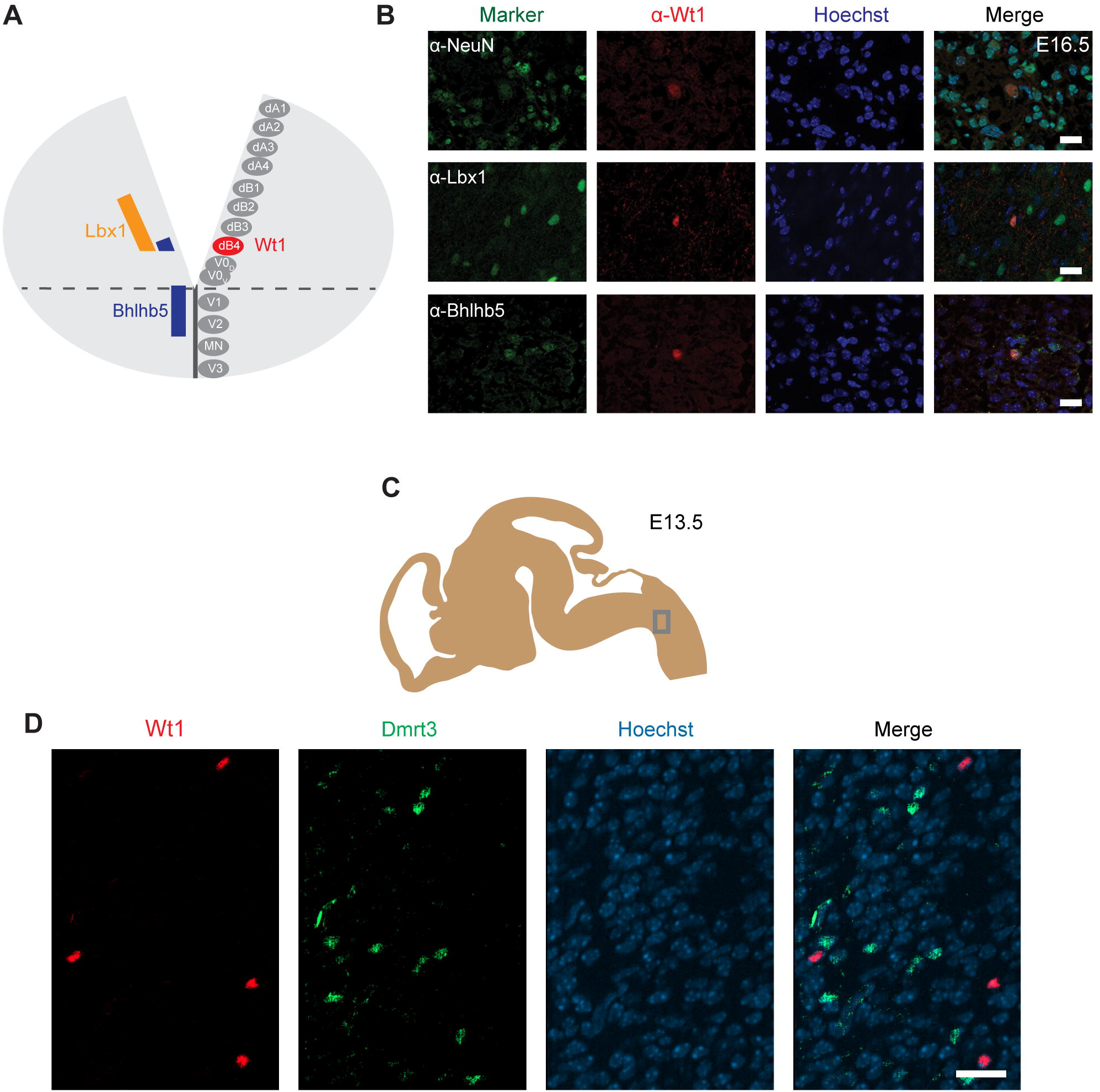
Wt1+ cells in the caudoventral medulla are dB4 neurons. **(A)** Schematic illustration of embryonic hindbrain. Neurons in the hindbrain arise from various progenitor domains surrounding the central channel: 8 dorsal neuron domains (dA1 till dB4), four ventral neuron domains (V0 till V3) and one motor neuron domain (MN). The markers of the various neurons were used to determine the dB4 domain to be the one that Wt1+ neurons (red) arise from in the hindbrain. **(B)** Hindbrain sections from E16.5 embryonic mice used for immunofluorescence analyses of Wt1+ neurons with the neuronal marker NeuN and particular markers (Lbx1; Bhlhb5) present in dB4 and adjacent neuron populations. Overlapping localization of Wt1, Lbx1 and Bhlhb5 reveals dB4 origin for Wt1+ neurons. Scale bar: 20 µm. **(C)** The scheme shows the area from which the image in D was taken. Developmental stage: E13.5. **(D)** Immunofluorescence analyses showed Dmrt3+ cells (green) to be present in the ventral hindbrain in proximity to Wt1+ neurons (red). No co-localization of Dmrt3 and Wt1 was observed. Scale bars: 20 µm

A subpopulation of dI6 neurons in the spinal cord is characterized by the presence of the transcription factor Dmrt3. This subpopulation overlaps, in part, with Wt1^+^ cells giving rise to a small Dmrt3/Wt1 double-positive population of dI6 neurons (Andersson et al., 2012). Dmrt3 has been suggested but not yet shown to occur in dB4 neurons as well. Immunofluorescence analyses revealed that Dmrt3+ cells are indeed present in the ventral hindbrain and that they are situated in close vicinity to the Wt1+ neurons (Fig. 3C). However, no co-localization was observed between Wt1 and Dmrt3 (Fig. 3D).

### Alterations in neuron composition upon Wt1 inactivation

Having determined the spatial and temporal distribution of the Wt1+ neurons, we aimed to apply the conditional Wt1 knockout mouse model *Lbx1-ki-Cre;Wt1^fl/fl^* to reveal possible functions of Wt1 in the dB4 cells. To verify the conditional *Wt1* deletion, embryos at stage E16.5 were used for immunohistochemical analyses of Wt1 presence in the ventral medulla. *Lbx1-ki-Cre;Wt1^fl/fl^* embryos did not show any Wt1+ neurons in the ventral region of the medulla where the cells occur in control animals (Fig. 3A). The Wt1+ cells in the AP and the Wt1+ cells in the choroid plexus were not affected by the conditional *Wt1* knockout in *Lbx1-ki-Cre;Wt1^fl/fl^* animals. This shows that neither of those populations expresses *Lbx1* and thus are not dB4 neurons. Thus, this line allows specific deletion of *Wt1* in the neurons of the ventral medulla already at embryonic stage.

Having observed alterations in the neuron composition of the developing spinal cord upon deletion of *Wt1* (Schnerwitzki et al., 2018), we intended to verify if the extent of several neuronal subtypes is changed in the hindbrain of *Wt1* knockout embryos. We focused on those ventral neuron subtypes that we had found to be altered in the spinal cord, namely Evx1+ V0 neurons and Chx10+ V2a neurons.

Due to the unequal distribution of Wt1, Chx10 and Evx1+ cells in the hindbrain, their respective cell numbers depend strongly on the orientation (coronal or sagittal) and the spatial level of the section. Therefore, cell numbers were normalized to a defined area (908 µm × 908 µm) in sagittal sections (Fig. 4B). As shown before, Wt1+ neurons are distributed ventral of this area in a stripe-like fashion (Fig. 4C). Chx10 and Evx1+ cells occur in close proximity to Wt1+ cells in the ventral hindbrain (Fig. 4 D, E). The population labelled by Chx10 extends further rostral and dorsal than the Wt1 population (Fig. 4D). Besides, the Evx1+ cells are enriched in the ventral area but show an additional diffuse distribution over the whole hindbrain (Fig. 4E). Therefore, all numbers are based on cells in the vicinity to both columns where the Wt1+ neurons occur.

**Fig. 4.**
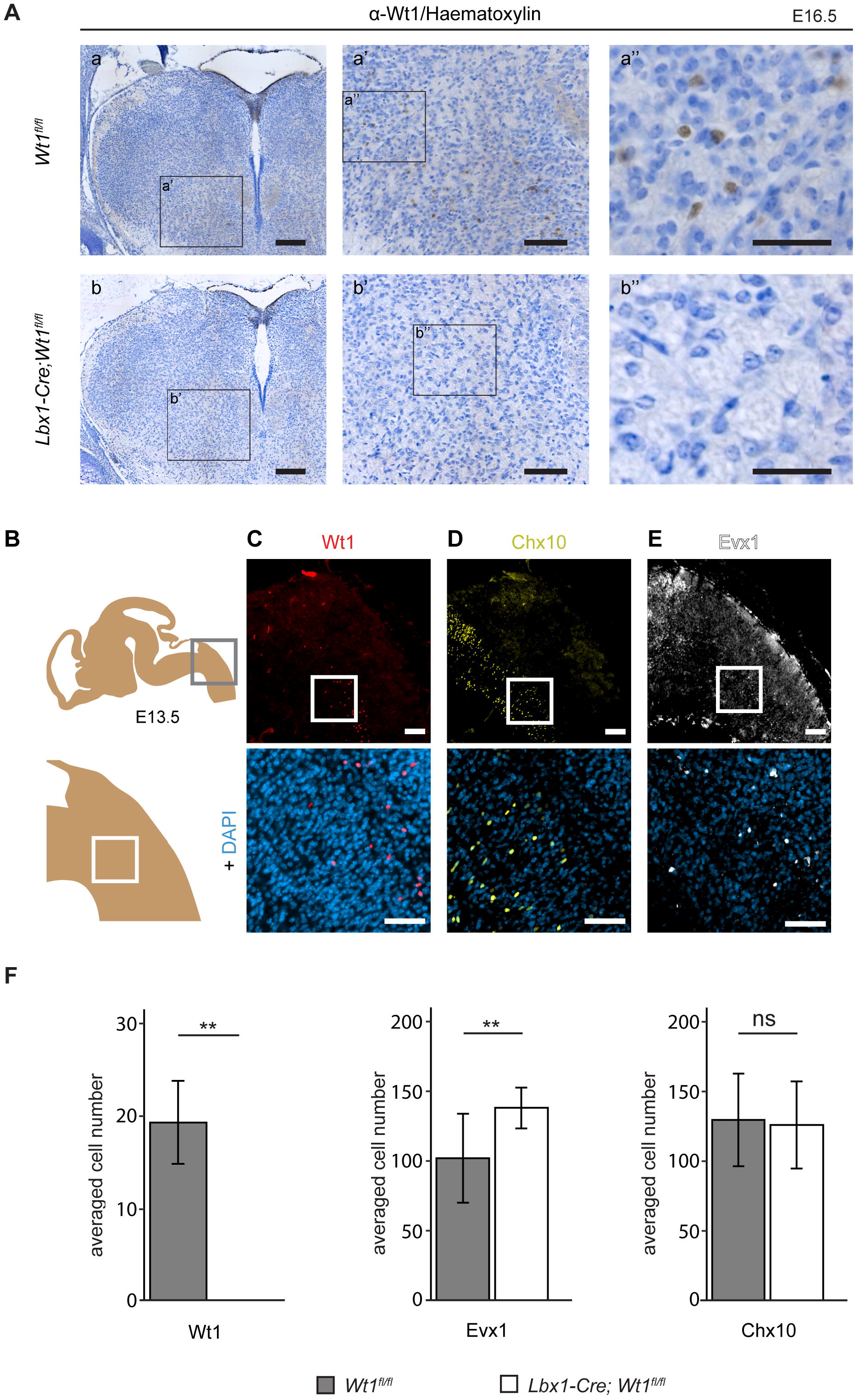
Neuron composition in conditional *Wt1* knockout embryos. **(A)** Immunohistochemical analyses of Wt1 on hindbrain sections from *Lbx1-ki-Cre;Wt1^fl/fl^* embryos (E16.5) and control animals were used to verify deletion of *Wt1* in the ventral medulla of *Lbx1-ki-Cre;Wt1^fl/fl^* animals. In order to assure that the same hindbrain region of *Lbx1-ki-Cre;Wt1^fl/fl^* and control embryos was examined, the non-neuronal Wt1+ cells found in the V-shaped AP were used as landmarks. Higher magnification images (a’; a’’ and b’; b’’) show that Wt1 is not detectable in the ventral medulla of *Lbx1-ki-Cre;Wt1^fl/fl^* compared to control animals. Orientation of the images: dorsal-up and ventral-down. Scale bar left column: 100 μm; middle column: 100 μm; right column: 50 μm. **(B)** Upper panel: area (grey frame) in which the Wt1, Chx10 and Evx1+ cells of Wt1 knockout and control mice were counted. Lower panel: area (white frame) that was magnified to visualize the cells in **C, D** and **E:** Immunofluorescence staining of sagittal embryonic hindbrain sections (E13.5) show the area harboring the Wt1, Evx1 and Chx10+ cells that were used for quantification (upper panel). Higher magnification together with Hoechst staining was used to confirm nuclear localization of the respective staining for Wt1, Evx1 and Chx10+ cells (lower panel). Wt1+ neurons are distributed in the ventral part (C). Evx1 labelled cells are diffusely distributed over the hindbrain (D). The population of Chx10+ cells extends further rostral and dorsal (E). Scale bars: 100 µm. **(F)** Quantification of the average cell number of Wt1, Chx10 and Evx1+ cells in hindbrain of conditional Wt1 knockout embryos (*Lbx1-cre;Wt1^fl/fl^*) reveals an absence of Wt1+ neurons compared to control individuals. The number of Chx10+ cells remains constant upon *Wt1* knockout but the number of cells labelled by Evx1 increases. n= 3; Linear Mixed Model: p < 0.01 (**), not significant (ns).

The quantitative analysis of the cell number in that defined area confirms the absence of Wt1+ cells in the ventral region of E13.5 *Lbx1-ki-Cre;Wt1^fl/fl^* embryos (Fig. 4F). Since the Wt1+ cells of the AP turned out not to be dB4 neurons, they were also not considered for counting. The determination of the number of Evx1 and Chx10+ cells revealed a significant increase in the amount of Evx1+ cells in the respective area upon deletion of *Wt1* while the number of Chx10+ cells did not change at developmental stage E13.5. This decline in the number of Wt1-positive dB4 neurons and the concomitant increase in the amount of Evx1+ cells might point to a change in the developmental fate from dB4 neurons into V0 neurons upon loss of Wt1.

### Wt1+ dB4 neurons are crucial for respiration

In order to analyze the putative involvement of the Wt1+ dB4 neurons in respiration, adult *Lbx1-ki-Cre;Wt1^fl/fl^* were examined for their respiration rate. The latter was determined by recording X-ray radiographs of mice at rest in order to count the number of inspirations by the movement of the diaphragm (Suppl. Vid. 2; Fig. 5A). The mean respiration rate was calculated as the number of inspirations per second (Fig. 5B). For *Lbx1-ki-Cre;Wt1^fl/fl^* mice, the mean respiration rate was significantly decreased compared to control animals. These changes in respiration rate were observed for both sexes but the effect was more pronounced in females. Thus, the alterations that we observed in conditional *Lbx1-ki-Cre;Wt1^fl/fl^* mouse embryos, namely the loss of Wt1 in dB4 neurons and the concomitant increase in the amount of Evx1+ neurons manifests itself as a change in respiration rate in adult mice.

**Fig. 5.**
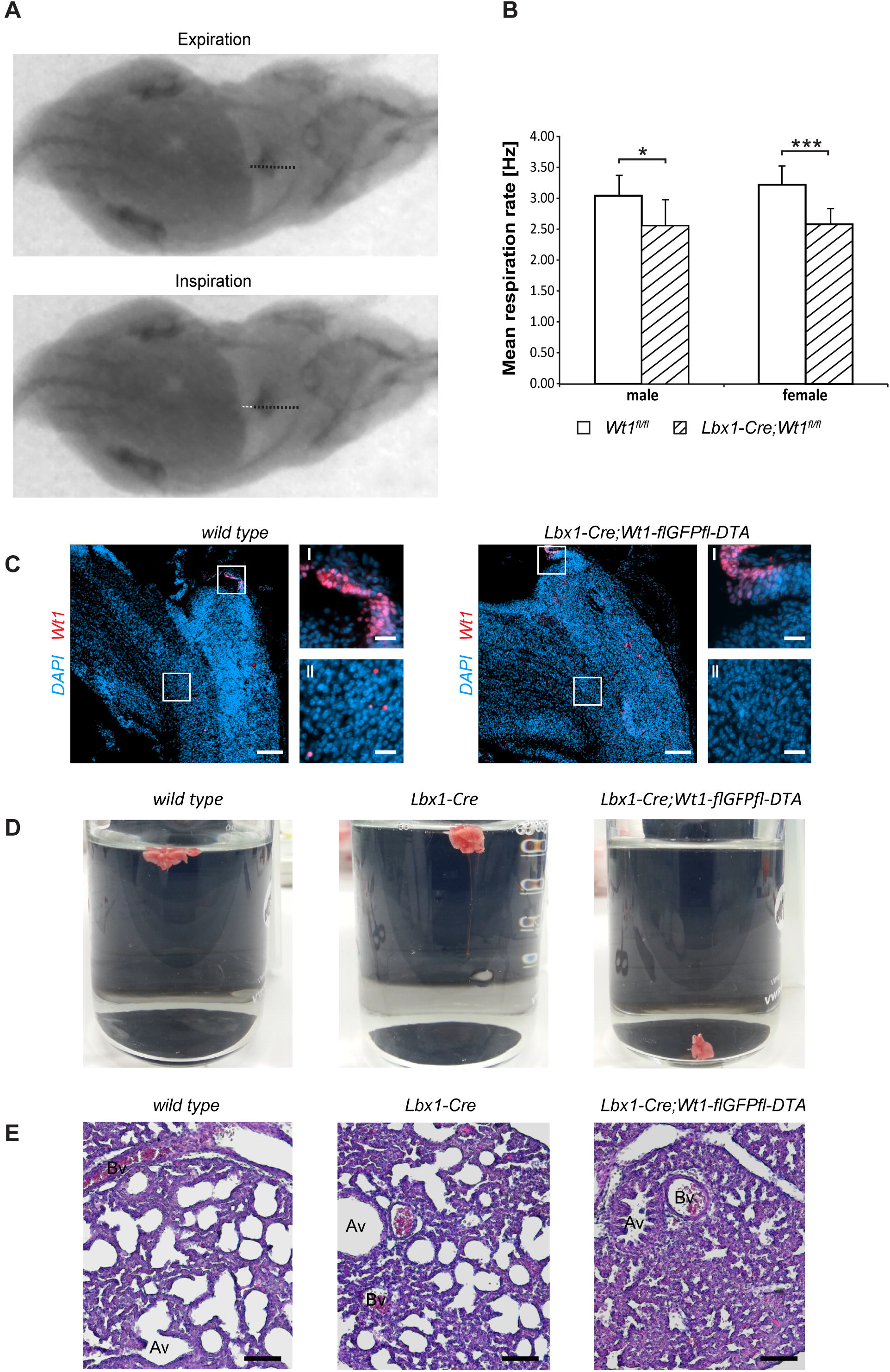
*Wt1*+ dB4 neurons are crucial for respiration. **(A)** Respiration rate was determined by recording X-ray radiographs of mice at rest. The movement of the diaphragm over time was analyzed (white dotted line in bottom image) and the mean respiration rate was calculated as the number of inspirations per second. Orientation of mice: ventral view with posterior on the left and anterior on the right. **(B)** The mean respiration rate was determined for *Lbx1-ki-Cre;Wt1^fl/fl^* mice and control animals. In both male and female *Lbx1-ki-Cre;Wt1^fl/fl^* mice the respiration rate is significantly decreased compared to controls. For both sexes n=10 *Wt1^fl/fl^* control; n=10 *Lbx1-ki-Cre-Cre;Wt1^fl/fl^*. Data are expressed as mean ± SD. Significance level: *** P<0.001; * P<0.05 (according to Student’s *t*-test). **(C)** Immunohistochemical analyses of Wt1 on hindbrain sections from *Lbx1-ki-Cre;Wt1-flGFPfl-DTA* embryos (E13.5) and wild type animals verify ablation of *Wt1*+ cells in the ventral medulla of *Lbx1-ki-Cre;Wt1-flGFPfl-DTA* embryos. In order to assure that the same hindbrain region of *Lbx1-ki-Cre;Wt1-flGFPfl-DTA* and control embryos was examined, the non-neuronal Wt1+ cells found in the AP were used as landmarks (magnification I). Higher magnification images (magnification II) show that Wt1+ cells are not detectable in the ventral medulla of *Lbx1-ki-Cre;Wt1-flGFPfl-DTA* compared to wild type animals. Scale bars overview: 100 µm. Scale bars magnification: 20 µm. **(D)** Neonates of *Lbx1-ki-Cre;Wt1-flGFPfl-DTA* exhibit respiratory failure. Lung hydrostatic tests show sinking of lungs from *Lbx1-ki-Cre;Wt1-flGFPfl-DTA* neonates as they have never inflated their lungs properly after birth. This is in contrast to lungs from control wild type and *Lbx1-ki-Cre* animals whose lungs are floating due to inspiration and inflation of lungs with air. **(E)** Histological sections of lungs from control animals and *Lbx1-ki-Cre;Wt1-flGFPfl-DTA* neonates were stained with eosin and haematoxylin. The alveoli of control animals are inflated with air. In *Lbx1-ki-Cre;Wt1-flGFPfl-DTA* neonates, the alveoli have never inflated after birth. (Av – alveolus; Bv – blood vessel).

We next wanted to examine the role of the Wt1-positive dB4 neurons as such. As reported earlier (Schnerwitzki et al., 2018), *Lbx1-Cre;Wt1-GFP-DTA* mice show ablation of Wt1- and Lbx1-co-expressing cells in the spinal cord due to the diphtheria toxin subunit A (*DTA*) which is expressed from the endogenous *Wt1* locus after Cre-mediated excision of a *GFP* cassette harboring a translational STOP codon. An ablation of Wt1+ cells was also detected in the embryonic hindbrain. As for the conditional knockout of Wt1, only the Wt1+ dB4 neurons in the ventral medulla were effected whereas the Wt1+ cells in the AP and the choroid plexus could still be observed (Fig. 5C). When monitoring *Lbx1-ki-Cre;Wt1-flGFPfl-DTA* animals at birth, they were found to be vital as assessed by body movements after non-noxious stimuli of the skin. However, *Lbx1-ki-Cre;Wt1-flGFPfl-DTA* neonates did not exhibit gasping after birth, which would then turn into abdominal breathing, as seen for control littermates. *Lbx1-ki-Cre;Wt1-flGFPfl-DTA* did not show contraction of the abdominal wall and became cyanotic. Eventually, they did not respond to stimuli anymore and died.

To verify the phenotype of *Lbx1-ki-Cre;Wt1-flGFPfl-DTA,* which is the incapability of neonates to initiate proper respiration, lung hydrostatic tests and histological analyses of the lung were performed. In wild type animals, the lung inflates during the first breaths (Fig. 5D). When the lung is transferred into water, it floats due to the inflation with air. This effect was also seen with lungs from *Lbx1-ki-Cre* controls. Newborn *Lbx1-ki-Cre;Wt1-flGFPfl-DTA* animals, however, did not breathe and never inflated their lungs properly. As a result their lungs sank.

The missing inflation of the lung was also detected histologically (Fig. 5E). The alveoli of wild type and *Lbx1-ki-Cre* animals were inflated seen by their round shape. Alveoli from *Lbx1-ki-Cre;Wt1-flGFPfl-DTA* neonates, however, were uninflated. These morphological alterations confirm that *Lbx1-ki-Cre;Wt1-flGFPfl-DTA* neonates are impaired in developing a proper respiration at birth.

These data show that the Wt1+ dB4 neurons in the medulla are involved in regulating respiration and that deletion of *Wt1* leads to a decreased respiration rate. Moreover, the existence of these Wt1+ dB4 neurons is a prerequisite for newborn animals to initiate respiration as ablation of these cells leads to death at birth.

## Discussion

In the present study, we show that *Wt1* is expressed in dB4 neurons during mouse embryonic and postnatal development. These Wt1+ neurons are located in one major and one minor column in each hemisphere of the medulla. If *Wt1* is deleted in dB4 cells at an embryonic stage, the neuronal composition of the hindbrain is altered. These changes also have late effects in that the respiration rate of adult mice with *Wt1* inactivation in dB4 neurons is decreased. More severe than *Wt1* deletion is the loss of the Wt1+ dB4 neurons in the hindbrain as these cells are indispensable for breathing.

The *Lbx1-ki-Cre;Wt1-flGFPfl-DTA* neonates show ablation of cells expressing both *Wt1* and *Lbx1*. Surprisingly, the newborns die immediately after birth provoking the idea that cells are depleted which are essential for life. Histological analyses showed impairments of *Lbx1-ki-Cre;Wt1-flGFPfl-DTA* neonates to develop a proper respiration at birth pointing to the fact that cells expressing both *Wt1* and *Lbx1* are involved in establishing respiration. Besides a shared expression of *Lbx1* and *Wt1* in the dB4 neurons of the medulla, both genes have been reported to be co-expressed in dI6 neurons of the spinal cord (Haque et al., 2018; Schnerwitzki et al., 2018) as well as in cells of the heart and diaphragm (Chao et al., 2011; Dingemann et al., 2011). Of those organs, the diaphragm is essential for respiration, too. The question may arise whether the inability to initiate respiration in *Lbx1-ki-Cre;Wt1-flGFPfl-DTA* neonates might be also due to ablation of cells in the diaphragm. Although *Lbx1* and *Wt1* are expressed in the diaphragm, the expression pattern of both genes is mutually exclusive. *Lbx1* is expressed in myogenic cells (Chao et al., 2011), whereas expression of *Wt1* was described in non-muscular components of the developing pleural mesothelium (Dingemann et al., 2011). That points to the fact that ablation of the Lbx1 and Wt1+ dB4 neurons in the medulla seems to be causative for the neonatal lethality.

A homology between Wt1+ dB4 neurons in the hindbrain and dI6 neurons of the spinal cord has been proposed on the basis of certain markers that occur in both neuronal types during development (Hernandez-Miranda LR, 2016). We were able to confirm this proposal not only on the level of the occurrence of common transcription factors but also on the behavioral level. Thus, tissue-specific knockout of *Wt1* in dB4 neurons using the *Lbx1-ki-Cre;Wt1^fl/fl^* mouse model showed functional homologies to the dI6 neurons in the spinal cord. Those animals are vital but show a decreased respiration rate at rest comparable to the decline in stride frequency in the animals with *Wt1* deletion in dI6 neurons (Schnerwitzki et al., 2018). This decreased respiration rate resembles the slower respiratory rhythm observed in constitutive, homozygous *Lbx1* knockout embryos. It was reported that *Lbx1* expression in particular neurons of the medulla is necessary to avoid respiratory defects (Pagliardini et al., 2008). Here, a proper rhythm was shown to be generated by the pre-Bötzinger cells in these embryos at E18.5. This rhythm, however, is not transmitted properly to the motor neurons in the phrenic nucleus of the spinal cord. As a consequence, these embryos die at birth due to respiratory failure. That already the deletion of *Lbx1* leads to death of neonates might be explained by the fact that more neurons in the hindbrain express *Lbx1* than *Wt1*. The knockout of *Lbx1* thus affects more neurons involved in various motoric and sensory tasks. Heterozygous *Lbx1* knockout animals are not affected. They are vital and exhibit normal respiration. The homology in development between Wt1+ dB4 neurons and dI6 neurons becomes also apparent on the level of the cellular consequences of *Wt1* deletion. As our work shows, the number of Evx1+ cells increases upon *Wt1* knockout already at developmental stage E13.5. The Evx1+ neurons in the hindbrain derive from the V0 progenitor domain (Bouvier et al., 2010), which give rise especially to excitatory premotor neurons in the rVRG (Wu et al., 2017). That increase in the number of Evx1+ V0 neurons was also observed in embryonic spinal cord where a fate change from Wt1+ neurons to V0-like neurons has been suggested when *Wt1* is deleted (Schnerwitzki et al., 2018).

Another neuronal population whose number was found to be altered in the spinal cord due to a knockout of *Wt1* is the population of Chx10+ V2a neurons. In the hindbrain, Chx10+ cells have been reported to be situated in the ventral respiratory column without further characterization of their function or a putative involvement in respiration (Brown et al., 2014). Contrary to the spinal cord, their number is not changed in *Wt1* knockout embryos at stage E13.5. This might be due to a different functionality of these cells compared to the spinal cord or due to the time point during development when this decrease in the cell number manifests. Since the number of Chx10+ neurons in spinal cord does not decrease until stage E16.5, the number of those cells in the hindbrain might change later than the investigated stage E13.5.

In general, the behavioral and cellular studies reveal that Wt1+ neurons in the hindbrain are involved in neuronal CPG circuits responsible for rhythmic movements of the respiratory musculature as we have reported for their homologs in the spinal cord have been reported. In both regions of the CNS these cells seem to fulfil homologous functions as in both cases *Wt1* deletion leads to phenotypes associated with an altered neuronal composition and a slowed rhythm of movements.

Since the deletion of *Wt1* as well as the ablation of the Wt1+ dB4 neurons are both associated with defects in respiration and Wt1+ neurons are to be found in columns in the caudal region of the medulla, the Wt1+ cells are likely to belong to the caudal portion of the ventral respiratory column in particular the cVRG (Fig. 1A). The cVRG is not well characterized in terms of their spatial dimension and cellular composition. However, it harbors some cells described for their involvement in respiration (Smith et al., 2013). Premotor neurons and third order neurons of motor neurons in the phrenic nucleus are located in the cVRG (Dobbins and Feldman, 1994). One type of those neurons in the cVRG are the so called early expiratory neurons firing in a declining manner (E-DEC neurons). They are inhibitory neurons that participate in the signaling necessary for expiration (Ezure et al., 2003). The Wt1+ dI6 neurons of the spinal cord are inhibitory neurons, too. (Haque et al., 2018). Since there seems to be a strong homology between development of Wt1+ neurons in the hindbrain and spinal cord, Wt1+ dB4 neurons of the hindbrain are also presumably inhibitory and thus might be at least a fraction of E-DEC neurons. To test this hypothesis, the electrophysiological properties of the Wt1+ dB4 neurons have to be determined applying the *Wt1^GFP^* mouse model which we have already used to label the Wt1+ neurons in the hindbrain.

Taken together, the reports from the *Lbx1* knockout and the findings shown in our study suggest that the dB4 neurons in the cVRG, which express both *Lbx1* and *Wt1,* are part of the respiratory CPG and are crucial for breathing. As suggested for cVRG neurons (Smith et al., 2013), they seem to transmit the rhythmic signals generated in the pre-Bötzinger complex to the motor neurons in the phrenic nucleus that innervate the muscles of the diaphragm. Deletion of either *Wt1* or *Lbx1* leads to changes in the transmission of these vital signals that are associated with a decreased respiratory rhythm. Ablation of Wt1+ dB4 neurons seems to “cut the wire” between the rhythm generating pre-Bötzinger complex and motor neurons. To prove the putative transmitting position of the Wt1+ dB4 neurons, it will be necessary to verify the proper function of the rhythm generating pre-Bötzinger complex in *Lbx1-ki-Cre;Wt1-flGFPfl-DTA* embryos at birth by using calcium imaging and electrophysiological recordings. Additionally, a putative loss of motor neuron output of the phrenic nerve, which innervates the muscles of the diaphragm, could be detected by recording currents at the ventral roots of the cervical level C3 – C5 of the spinal cord.

In sum the results obtained from our work confirms not only the homology between the development and function of dB4 and dI6 neurons in the hindbrain and spinal cord but also reveal the so far undescribed necessity for Wt1+ dB4 neurons for respiration and thereby their indispensability for life.

## Materials and Methods

### Mouse husbandry

All mice were bred and maintained in the Animal Facility of the Leibniz Institute on Aging – Fritz Lipmann Institute (FLI), Jena, Germany, according to the rules of the German Animal Welfare Law (Animal licences: J-SHK-2684-04-08-01/14, J-SHK-2684-04-08-02/14, TG/J-0002858/A, TG/J-0003616/A, TG/J-0003681/A, 22-2684-04-03-004/14, 22-2684-04-03-049/16). Sex- and age-matched mice were used. Animals were housed under specific pathogen-free conditions (SPF), maintained on a 12 hour light/dark cycle and fed with mouse chow and tap water *ad libitum*. *Wt1^fl/fl^* mice were maintained on a mixed C57B6/J x 129/Sv strain. *Wt1^GFP^* mice (Hosen et al., 2007) were maintained on a C57B6/J strain. Conditional *Wt1* knockout mice were generated by breeding *Wt1^fl/fl^* (Gebeshuber et al., 2013) to *Lbx1-Cre;Wt1^fl/fl^* mice (Sieber et al., 2007). To generate mice with Wt1 ablated cells, *Wt1-flGFPfl-DTA* mice (Schnerwitzki et al., 2018) were bred with *Lbx1-Cre* mice. Control mice were sex- and age-matched littermates (wild type or *Wt1^fl/fl^*). For plug mating analysis, females of specific genotypes were housed with males of specific genotypes and were checked every morning for the presence of a plug. For embryo analysis, pregnant mice were sacrificed by CO_2_ inhalation at specific time points during embryo development and embryos were dissected. Typically, female mice between 2 and 6 months were used.

### Analysis of respiration rate of adult mice at rest

10 animals per sex and genotype (*Lbx1-ki-Cre;Wt1^fl/fl^* and *Wt1^fl/fl^* control) were used. We recorded the respiration rate of adult mice using high-resolution X-ray fluoroscopy (biplanar C-arm fluoroscope Neurostar, Siemens AG, Erlangen, Germany). The X-ray system operates with high-speed cameras and a maximum spatial resolution of 1536 dpi × 1024 dpi. A frame frequency of 500 Hz was used. A normal-light camera operating at the same frequency and synchronized to the X-ray fluoroscope was used to document the entire trial from the lateral perspective. Respiration rate was determined by recording the movement of the diaphragm of mice at rest. The mean respiration rate was calculated for each mouse as the number of inspirations per second. Group means were calculated from the means of the 10 animals. Student’s t-test was computed to determine significance of the differences between the means of *Wt1^fl/fl^* and *Lbx1-ki-Cre;Wt1^fl/fl^* animals.

### Lung hydrostatic test

Lung hydrostatic test was performed with lungs from *Lbx1-Cre;Wt1-flGFPfl-DTA* neonates to determine whether lungs had been inflated after birth. Therefore, the entire lung was removed from the thorax and subsequently transferred into water. Floating of the lung pointed to proper inflation with air. Sinking of the lung implied absence of air in the lungs.

### Haematoxylin and eosin staining

The tissue was processed using an automated slide stainer (Leica, Wetzlar/Germany), with the following program: paraffin sections were deparaffinized with xylene 2x for 10 min, followed by a series of rehydration steps using a decreasing gradient of ethanol. The tissue was stained in eosin for 2 min followed by washing in water, further followed by haematoxylin staining for 1.5 min. Stained sections were dehydrated using an increasing gradient of ethanol and cleared in xylene. Finally, slides were mounted using xylene based mounting medium.

### Immunohistochemistry staining on sections

Paraffin sections were deparaffinized as described above. To retrieve antigens, slides were incubated in sub boiling sodium citrate buffer for 30 minutes. During the incubation, the temperature was continuously checked to be between 98°C – 100°C. After antigen retrieval, slides were cooled down to RT and subsequently washed with H_2_O. To permeabilize the tissue and saturate endogenous peroxidases, the slides were incubated in methanol with 0.3% hydrogen peroxide for 20 min. After washing with PBS, the sections were incubated with the primary antibody (Wt1 1:50, Agilent - Dako, Santa Clara, California, USA) diluted in PBS at 4°C overnight. The next day, the slides were washed with PBS and incubated with the polymeric horse radish peroxidase (HRP)-conjugated secondary antibody for 30 min. After washing with PBS, the slides were incubated with 3,3’-Diaminobenzidine (DAB) solution for 6 min. The staining intensity was checked with the microscope and finally stopped with water. A counterstaining with hematoxylin was performed and the slides were mounted using xylene based mounting medium.

### Immunofluorescence staining on sections

Embryonic and postnatal brains were dissected. They were either frozen unfixed after 15 min dehydration with 20% sucrose (in 50 % TissueTec/PBS) (post-fix) or fixed for 75 min in 4% paraformaldehyde in PBS (pre-fix). Pre-fixed tissue was cryo-protected in 10%, 20% and 30% sucrose (in PBS) before freezing in cryo-embedding medium (Neg-50 - Thermo Scientific, Kalamazoo, USA). Post- and pre-fix samples were sectioned (12 μm). Post-fixed samples were fixed for 10 min after sectioning and washed with 2% Tween in PBS (PBS-T). For pre-fixed samples, antigen retrieval was performed by incubation in sub boiling 10 mM sodium citrate buffer pH6.0 for 30 min. After blocking with 10% goat serum and 2% BSA in PBS-T (post-fix) or with 0.1% TritonX 100, 1% donkey serum or goat serum in PBS (prefix), sections were incubated with primary antibodies (in blocking solution) using the following dilutions: Chx10 1:100 (abcam, Cambridge, UK), Dmrt3 1:5000 (custom made (Andersson et al., 2012)), Evx1 1:1000 (Developmental Studies Hybridoma Bank, University of Iowa, Iowa City, IA), GFP 1:1000 (abcam, Cambridge, UKNeuN, 1:500 (Merck, Darmstadt, Germany), Phox2b 1:500 (gift from G. Fortins, Paris), Wt1 1:100 (Santa Cruz Biotechnology, Inc., Santa Cruz, California, USA), Wt1 1:1000 (abcam, Cambridge, UK). Secondary antibodies (Alexa Fluor secondary antibodies, Thermo Fisher Scientific, Waltham, Massachusetts) were applied according to species specificity of primary antibodies. Hoechst was used to stain nuclei.

### Whole-mount immunofluorescence staining and tissue clearing

For rehydration, samples were treated consecutively with 75%, 50% and 25% methanol. After washing wit PBS-T, antigen retrieval was performed as mentioned above. Afterwards, samples were washed with PBS-T and incubated with blocking solution consisting of 5% NGS, 1% DMSO and 0.5% Triton X 100 diluted in PBS. Whole-mount tissue was incubated with GFP primary antibodies (diluted 1:500 in blocking solution) at 4°C for at least 24 hours. Samples were washed meticulously with PBS-T for another 24 h before incubation with secondary antibodies for at least for 24 hours. After washing with PBS-T 3 times 2 hours, Hoechst (1:100) was added to the sample for overnight incubation before samples underwent a final washing with PBS-T as done before. For subsequent clearing of the tissue, the whole-mount samples were transferred into Sca*l*eA2 solution (4M Urea, 10%Glycerol, 0.1%Triton; pH 7.7) (Hama et al., 2011) and incubated at 4°C for 10 months. The level of clearing was examined every 2^nd^ month and the Sca*l*eA2 solution was replaced with fresh one.

### Imaging and picture processing

Fluorescent images were acquired with a Zeiss Axio Imager and a Zeiss Axio Observer Z1 equipped with an ApoTome slider for optical sectioning (Zeiss, Germany). Cleared and whole-mount stained specimen were imaged with a light sheet microscope (Lightsheet Z1, Zeiss, Germany) enabled for dual side illumination and equipped wih a 20x detection objective suitable for clearing methods with refractive index of 1.38 (CLR Plan-Apochromat Corr nd=1.38 VIS-IR, numerical aperture = 1.0, working distance = 5,6 mm). For image processing which consisted of dual side fusion, three-dimenional reconstruction and animation as well as brightness and contrast adjustment, ZEN software (black edition, Zeiss, Germany) was used.

### Statistical Analyses

Data are expressed as mean ± SD. Groups were compared using two-tailed two-sample Student’s *t*-test. All statistical analyses were done using Microsoft Excel (Microsoft Corporation, Redmond, USA) or Matlab (Mathworks, R2014b). Significance was determined as * = P <0.05, ** = P<0.01, *** = P<0.001.

## Acknowledgements

We thank Dagmar Kruspe, Rommy Peterson as well as Manuela Schmidt for technical assistance and Carmen Birchmeier (MDC, Berlin, Germany) for providing the *Lbx1-Cre* mouse line. D.S. received a scholarship from the Leibniz Graduate School on Ageing and Age-Related Diseases (LGSA). The FLI is a member of the Leibniz Association and is financially supported by the Federal Government of Germany and the State of Thuringia.

## Author contributions

D.S., C.E. initiated, coordinated and drove the project. D.S. organized and coordinated the mouse work. D.S. and C.H. performed the characterization of Wt1+ cells (special and temporal distribution; marker analyses). C.H. and B.P performed tissue clearing and light sheet microscopy. C.H. determined alterations in the neuronal subpopulations. D.S. performed behavioral analysis including analysis of the X-ray recordings for diaphragm movement. D.S., C.H, B.P. and C.E. wrote and revised the manuscript.

